# *Entamoeba histolytica* EHD1 is involved in mitosome-endosome contact

**DOI:** 10.1101/2022.01.16.476539

**Authors:** Herbert J. Santos, Yuki Hanadate, Kenichiro Imai, Haruo Watanabe, Tomoyoshi Nozaki

## Abstract

Inter-organellar crosstalk is often mediated by membrane contact sites (MCSs), which are zones where participating membranes come within a proximity of 30 nm. MCSs have been found in organelles including the endoplasmic reticulum, Golgi bodies, endosomes, and mitochondria. Despite its seeming ubiquity, reports of MCS involving mitochondrion-related organelles (MROs) present in a few anaerobic parasitic protozoa remain lacking. Entamoeba histolytica, the etiological agent of amoebiasis, possesses an MRO called mitosome. We previously discovered several Entamoeba-specific transmembrane mitosomal proteins (ETMPs) from in silico and cell biological analyses. One of them, ETMP1 (EHI_175060), was predicted to have one transmembrane domain and two coiled-coil regions, and was demonstrated to be mitosome membrane-integrated based on carbonate fractionation and immunoelectron microscopy (IEM) data. Immunoprecipitation analysis detected a candidate interacting partner, EH-domain containing protein (EHD1, EHI_105270). We expressed HA-tagged EHD1 in E. histolytica and subsequent immunofluorescence and IEM data indicated an unprecedented MCS between the mitosome and the endosome. Live imaging of GFP-EHD1 expressing strain demonstrated that EHD1 is involved in early endosome formation, and is observed in MCS between endosomes of various sizes. In vitro assay using recombinant His-EHD1 demonstrated ATPase activity. MCSs are involved in lipid transfer, ion homeostasis, and organelle dynamics. The serendipitous discovery of the ETMP1 interacting partner EHD1, led to the observation of the mitosome-endosome contact site in E. histolytica. It opened a new view of how the relic mitochondria of Entamoeba may likewise be involved in organelle crosstalk, a conserved feature of mitochondria and other organelles in general.

## Introduction

Membrane contact sites (MCSs) mediate communication and exchanges between membrane-bound compartments by the assembly of protein-protein or protein-lipid tethers which maintains distancing of 30 nm between interacting membranes. MCSs have been found in almost every pair of organelles (1), most of which involving the endoplasmic reticulum (ER) as its membrane spans a network that interacts with the plasma membrane, and other organellar membranes such as those of the Golgi apparatus, lysosomes, endosomes, lipid droplets, peroxisomes, and mitochondria (2). MCSs are also reported between other organelle pairs such including the peroxisomes and lipid droplets, and the mitochondria and vacuoles/endosomes/lysosomes, plasma membrane, lipid droplets, and peroxisomes respectively, notwithstanding the contact sites between the inner and outer mitochondrial membranes (3). These contact sites mostly harbor proteins involved in lipid metabolism and transport, making them hubs of lipid transfer between interacting membranes. However, other processes associated with MCSs have been reported and they include ion transport and homeostasis, apoptosis (1), and endosomal (4) and mitochondrial (5) fission, respectively.

We recently identified several key molecules that facilitate membrane contact sites in mitosomes, endosomes, and Golgi apparatus of *Entamoeba histolytica*, and aimed to study their roles and possible link to the parasitic nature of this amoeba*. E. histolytica* is an anaerobic unicellular protozoan parasite that infects the large intestine of humans, and causes amebiasis, a disease characterized by diarrhea, which is a major cause of death in children worldwide. Millions of individuals are infected; mostly in developing countries and the disease causes an estimated 100,000 deaths annually (6). Infection begins by the ingestion of infectious cysts, which are resistant to the acidic environment of the stomach, then cysts pass through the small intestine, and undergo excystation within the terminal ileum or colon, to the trophozoite stage. Trophozoites reproduce and encyst within the colon, where they are released in the environment via excretion of feces, thus completing one cycle of the fecal-oral transmission (Stanley, 2003). Invasive amoebic trophozoites destroy the muco-epithelial barrier of the host intestinal tract, inducing mucus overproduction, inflammation and dysentery. It could lead to the formation of extra-intestinal abscesses particularly in the liver (amoebic liver abscess), lungs, and brain. The virulence of this parasite is due to its ability to inflict damage to host cells and tissues, via parasite attachment to colonic epithelial cells, protease secretion to damage host cells and evade host immune response, and by ingestion of host cells via phagocytosis and trogocytosis. These processes involve intracellular trafficking and inter-organellar crosstalk, underscoring the role of vesicular transport and membrane contact sites (MCSs) not only in parasite biology but also in its virulence and pathogenesis.

Like other anaerobic parasitic protozoans, *E. histolytica* lacks canonical mitochondria, and instead has a highly divergent mitochondrion-related organelle (MRO) called mitosome. *Entamoeba* mitosomes contribute to parasitism (7) due to a compartmentalized sulfate activation pathway that leads to the formation of cholesteryl sulfate in the cytosol. This molecule induces stage conversion from the trophozoite to cyst form (8) a process that is essential for maintaining the parasite’s life cycle and mode of disease transmission. Apart from mitosomes, other amoebic organelles such as the ER and the Golgi apparatus also show less defined structural and compositional features compared with model organisms, however, they have been shown to contain orthologs of established endomembrane proteins (9–11). Our knowledge on MCSs in *Entamoeba* is extremely limited, with only the previously identified mitosomal membrane proteins ETMP30 (reported to interact with a Golgi-localized protein secretory pathway calcium ATPase) and EHI_099350 (reported to have dual localization in the mitosomes and the ER) (12–14). Other molecules that participate in tethering of these compartments and what roles these contact sites play in the cell are still unknown, making it imperative to dissect amoebic MCSs. These past observations point to the fact that mitosomes, although highly degenerate, are able to interact with other organelles in the cytoplasm, and such contacts often utilize lineage-specific membrane. Here we identified another mitosomal membrane protein, ETMP1 which interacts with a C-terminal Eps15 homology domain (EHD) containing protein, a member of the EHD protein superfamily involved in various endocytic processes. Studies on *E. histolytica* organelle interaction via the endocytic transport mechanism has accumulated over several decades, including proteins involved in cargo sorting regulation and endosome dynamics such as Rab GTPases (15) and endosomal sorting complex required for transport (ESCRT) proteins (16, 17). However, there are so far no reports on whether these molecules take part in MCS between endosomes and other parts of the cell.

## Results

### EHI_175060 is a lineage-specific mitosomal membrane protein

Our group previously searched for transmembrane domain-containing mitosomal proteins using a previously-developed prediction pipeline (12) in sought for proteins that could be lineage-specific receptors, channels, enzymes, and components of the import machinery or otherwise uncharacterized complexes on the outer and inner membranes of *Entamoeba histolytica* mitosomes. This resulted in the prediction of 25 protein candidates. Like the other 24 proteins in the list (12), EHI_175060 is unique to the lineage *Entamoeba*. Figure 1 shows a multiple sequence alignment of the protein sequence of EHI_175060 with that of its orthologs in other *Entamoeba* species (*E. moshkovskii*, *E. dispar*, and *E. nuttalli*). The protein has a predicted molecular mass of 29.5 kDa, and it contains two coiled-coil domains at the middle portion and a single transmembrane domain near the carboxyl terminus. It also lacks a predictable canonical N-terminal targeting sequence. Based on these characteristics, we name EHI_175060, as *Entamoeba*-specific transmembrane mitosomal protein 1 (ETMP1).

**Figure 1.**
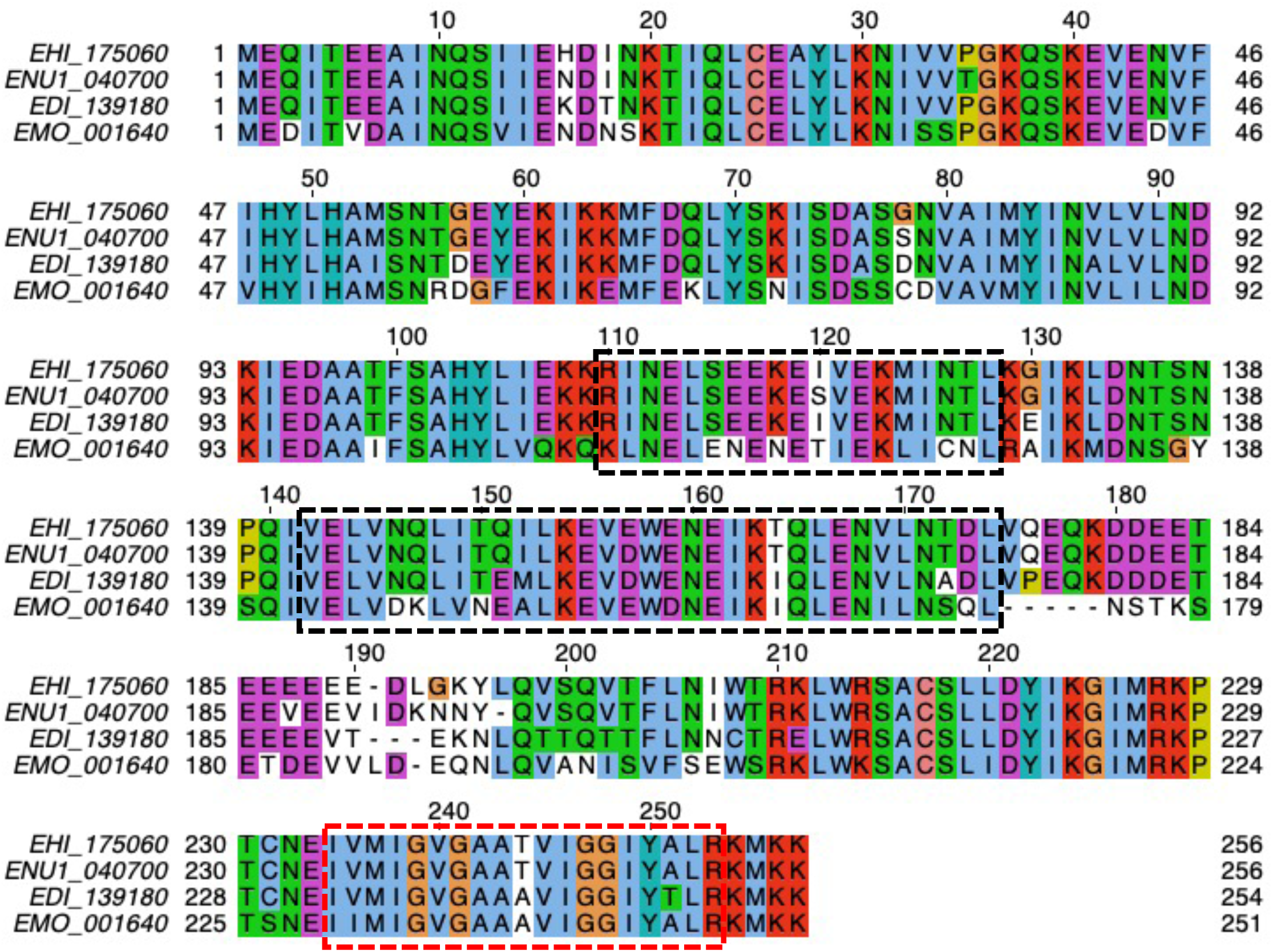
Multiple sequence alignment of ETMP1 orthologs in *Entamoeba*. Amino acid sequences of orthologs in *E. histolytica* (EHI_175060), *E. nutttalli* (ENU1_040700), *E. dispar* (EDI_139180), and *E. moshkovskii* (EMO_001640) were aligned using MAFFT (67), and displayed using Jalview (68). The hydrophobic, positively charged, negatively charged, polar, cysteine, glycine, proline, aromatic residues are indicated in blue, red, magenta, green, pink, orange, yellow, and cyan respectively. Dashed black boxes show predicted coiled coil domains by DeepCoil (64), while the dashed red box indicates the predicted transmembrane region by our TMD prediction tool (12).

### ETMP1 is localized to mitosomal membranes

To validate the predicted localization of ETMP1, we expressed amino terminus hemagglutinin (HA)-tagged fusion protein, HA-ETMP1, in amoebic trophozoites and confirmed protein expression by Western blot analysis. The anti-HA immunoblot showed a single band corresponding to the expected molecular mass of HA-ETMP1 (Figure 2A). We then analyzed the localization of the protein by immunofluorescence assay (IFA; Figure 2B). Co-staining of HA-ETMP1 expressing strain using anti-HA antibody and anti-adenosine-5’-phosphosulfate kinase (APSK; EHI_179080; a mitosomal matrix enzyme involved in sulfate activation) antiserum revealed good colocalization of the HA-tagged protein to mitosomes containing APSK. This is supported by the Pearson correlation R value which ranges between 0.31 to 0.59, suggesting that HA-ETMP1 is localized to mitosomes. Furthermore, we also performed Percoll-gradient fractionation of HA-ETMP1 homogenate and found that fractions containing HA-ETMP1 showed broad distribution in the first ultracentrifugation, suggesting some proteins are localized to the cytosol/lighter fractions. However, HA-ETMP1 also exists in the bottom fractions which overlap with those that contain chaperonin 60 (Cpn60; EHI_178570; a chaperone protein and canonical mitochondrial matrix marker). The co-fractionation of HA-ETMP1 to mitosomes was suggested by the anti-HA and anti-Cpn60 immunoblots of both the first and second ultracentrifugation respectively (Figure 2C). We also performed subcellular fractionation followed by carbonate treatment, to further assess the localization, as well as membrane integration of HA-ETMP1. The fractionation profile of HA-ETMP1 after immunoblot analysis (Figure 2D, upper panel) showed that it is present in both cytosolic and organelle fractions. Also, it was clearly demonstrated that the HA-ETMP1 contained in the organellar membrane-enriched fraction, is integrated to organellar membranes as it was retained in the particulate fraction after carbonate treatment, similar to MBOMP30-HA (Figure 2D, middle panel), a positive control for mitosomal membrane protein. These carbonate fractionation profiles contrast with that of the soluble mitosomal matrix protein marker Cpn60, as shown by the blot stained with anti-Cpn60 antiserum (Figure 2D, bottom panel).

**Figure 2.**
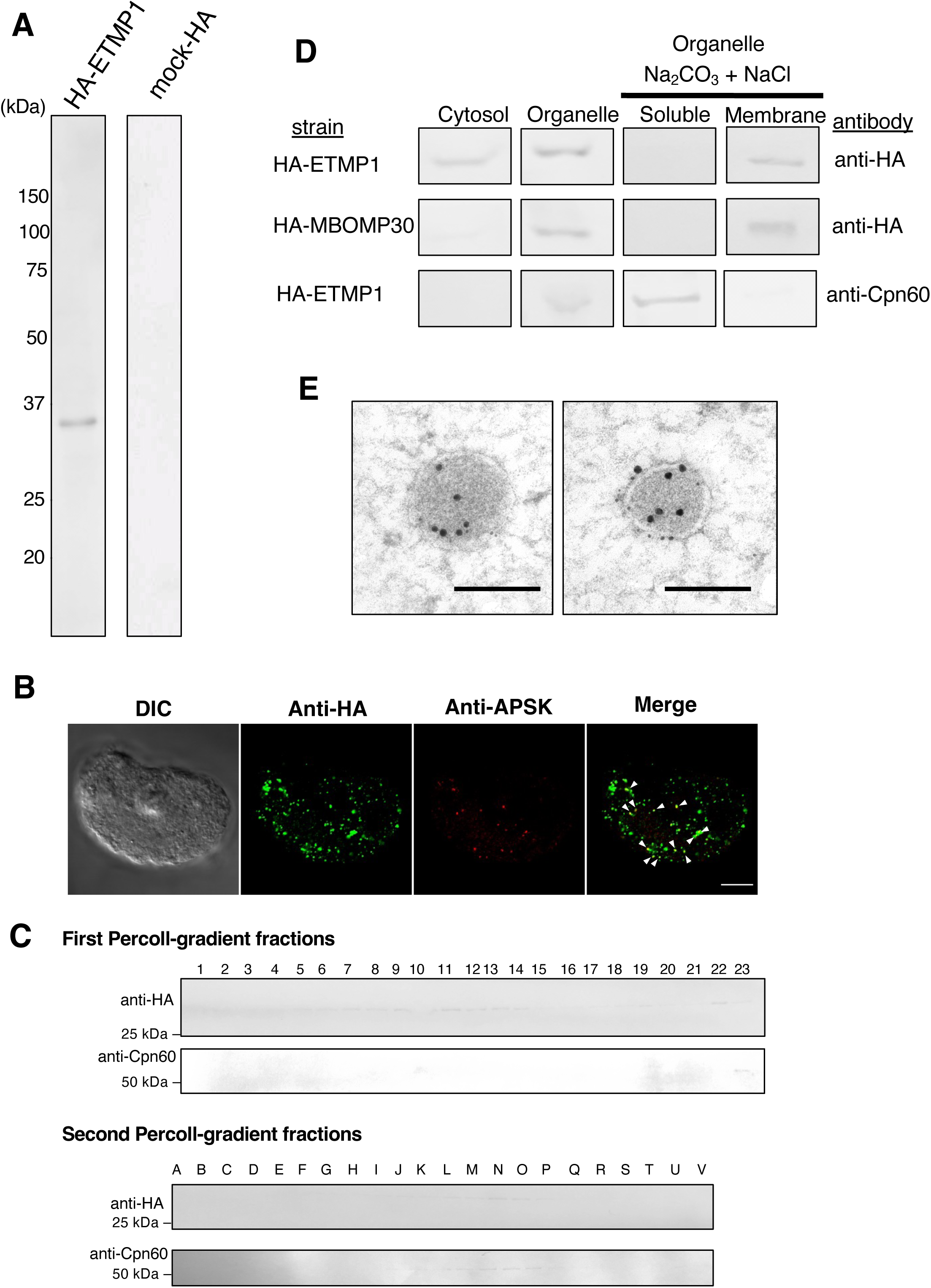
Expression and localization of HA-ETMP1 in *E. histolytica* trophozoites. (A) Approximately 30 µg protein from whole cell lysates of HA-ETMP1 and mock control (pEhEx-HA) strains were separated by SDS-PAGE and subjected to anti-HA immunoblot analysis. The 33 kDa band corresponds to the predicted molecular mass of HA-ETMP1 (white arrowhead). (b) Immunofluorescence analysis of HA-ETMP1-expressing trophozoites, double stained with anti-HA (green) and anti-APSK (red) respectively. Scale bar = 10 µm. (C) Fractionation of HA-ETMP1 by discontinuous Percoll-gradient ultracentrifugation. Homogenate of HA-ETMP1 was separated by density against a Percoll gradient. Approximately 15 µL of fractions collected from the first (1 to 22) and second (A to V) ultracentrifugation steps were separated by SDS-PAGE followed by immunoblot analysis with anti-HA and anti-Cpn60 antibodies respectively. (D) Anti-HA and anti-Cpn60 immunoblot profiles of subcellular fractionation including alkaline carbonate treated organelle-rich fractions of HA-ETMP1 and HA-MBOMP30 (mitosome membrane control) respectively. (E) Representative immunoelectron micrographs of 15 nm anti-APSK gold-labeled mitosomes of HA-ETMP1, co-stained with 5 nm anti-HA gold. Scale bar = 200 nm.

We also performed immunoelectron microscopy analysis, and the results indicated that HA-ETMP1 is localized to the mitosome membranes, as anti-HA gold particles are found along the periphery of the Cpn60-labeled mitosomes (Figure 2E). Particle distribution analysis of the gold-conjugated antibodies revealed a significant difference in the staining of mitosomes (368 ± 279/ μm^2^) compared to cytosol (22.3 ± 9.25/ μm^2^) by anti-HA gold. The distribution of the mitosomal marker APSK as detected by the gold anti-APSK particles was also significantly higher in mitosomes (192 ± 98.1/ μm^2^) than in the cytosol (0.984 ± 0.817/ μm^2^). Statistical significance in both datasets was analyzed using two-tailed Welch’s unequal variance t-test (n = 17, *p* < 0.0001). Overall, these data provide evidence of mitosomal membrane localization of ETMP1.

### ETMP1 is essential and its overexpression causes drastic growth defect

We made several attempts at silencing the *etmp1* gene by small-RNA transcriptional interference, all of which failed as the transformants did not survive drug selection, suggesting its essentiality to the parasite. We also observed a slower growth rate in HA-ETMP1 expressors compared with mock control. Analysis of growth kinetics of the two strains at varying concentrations of geneticin (G418) suggest a dose-dependent effect of drug concentration to the growth of amebic trophozoites (Figure 3A), and protein expression respectively (Figure 3B).

**Figure 3.**
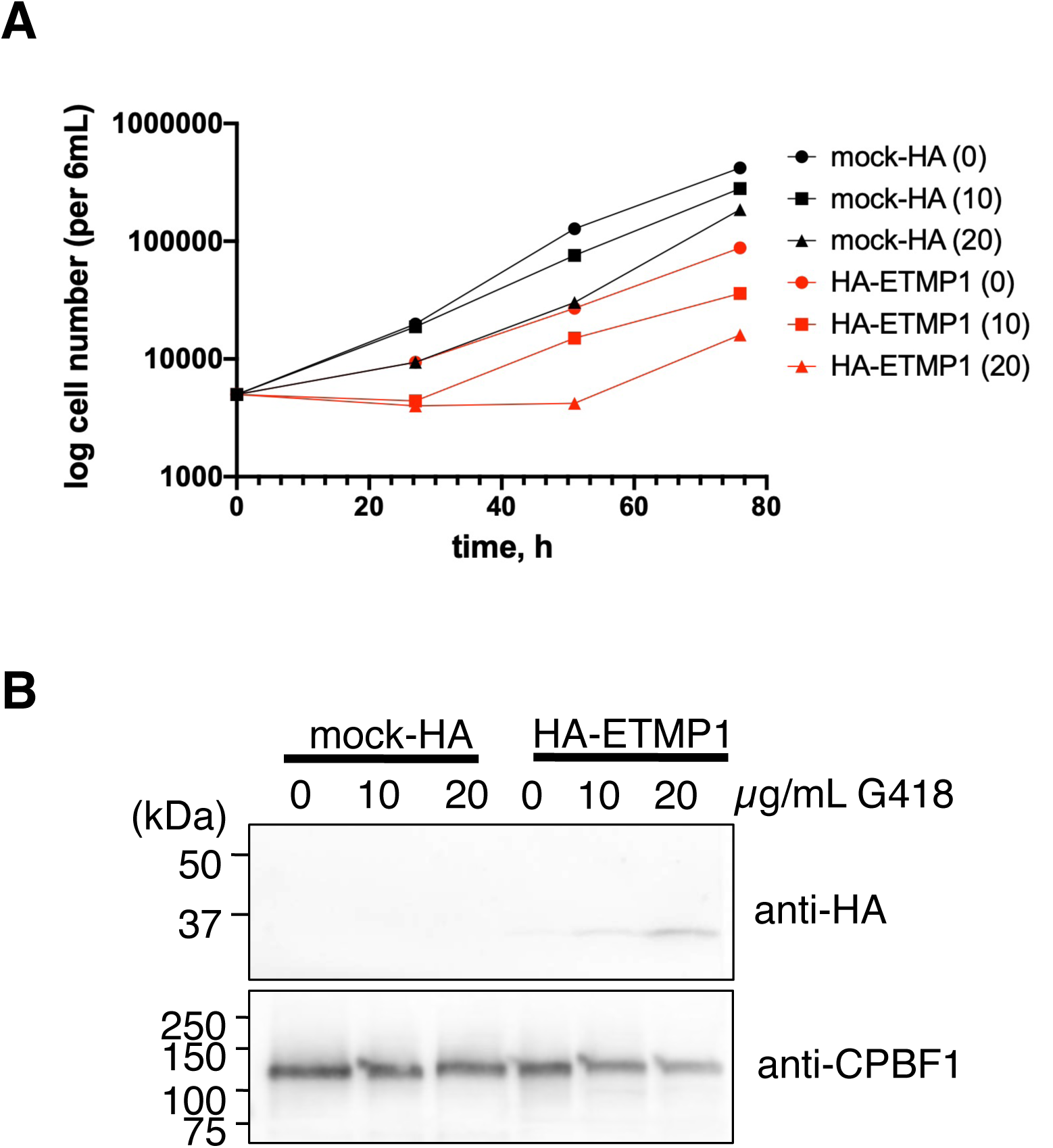
Growth curve of HA-ETMP1 and mock-HA strain. Cell numbers of ETMP1 (black line) and mock-HA strain (red line) cultivated in BI-S-33 medium containing 0, 10, and 20 µg/mL G418 respectively, were plotted against time (h). Western blot analysis of whole cell lysates of HA-ETMP1 and mock-HA grown in medium containing 0, 10, and 20 µg/mL G418, and harvested at various time points. Upper and lower panels show anti-HA and anti-CPBF1 (loading control) immunoblots respectively.

### ETMP1 interacts with EH-domain containing proteins

To shed light on the function of ETMP1, we next attempted to identify its interacting partner(s) by immunoprecipitation (IP). Anti-HA agarose beads were used to immunoprecipitate the bait protein together with its binding partner(s) from the organelle-rich fraction of HA-ETMP1-expressing and mock control strains respectively. Western blotting with anti-HA antibody confirmed successful binding to and elution from HA-ETMP1 with respect to the anti-HA beads (Figure 4A). Silver staining of the SDS-PAGE gel containing HA peptide-eluted fractions revealed a band corresponding to approximately 55 kDa that is uniquely precipitated in HA-ETMP1 (absent in the mock-HA control) (Figure 4B). Protein sequencing analysis by mass spectrometry followed by differential comparison of quantitative values (QVs), normalized with unweighted spectrum counts between HA-ETMP1 and mock-HA control identified interacting partners of ETMP1 (Figure 4C). A cutoff of QV >2.0 in the HA-ETMP1 over the mock-HA sample was used, yielded four candidates, three of which were exclusively detected in the eluted IP fraction of HA-ETMP1. Also, three of the four candidates were identified in the mitosome proteome that was previously published (18), namely, L-myo-inositol-1-phosphate synthase (EHI_070720), EH-domain (EHD) containing protein 1 (annotated as receptor mediated endocytosis protein; EHI_105270) and its close homolog, EHD2 (EHI_152680).

**Figure 4.**
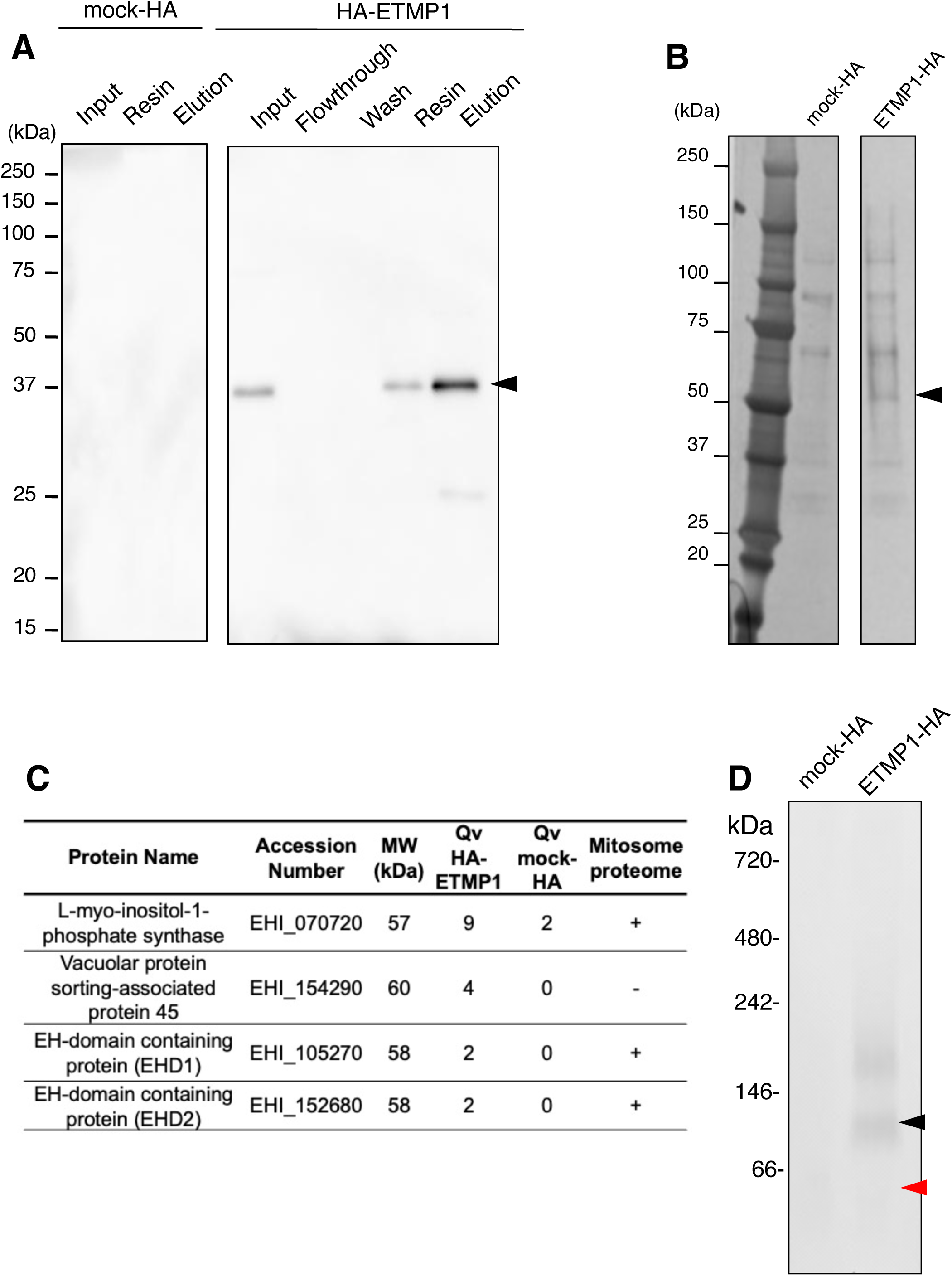
Anti-HA beads immunoprecipitation (IP) of mock-HA and HA-ETMP1 strains. (A) Western blot analysis using anti-HA antibody of the cell lysates and various IP fractions of HA-mock-HA (left) and ETMP1 (right) respectively. A black arrowhead indicates the position of HA-tagged ETMP1 (33 kDa). (B) Silver stained-SDS-PAGE gel of IP eluates of mock-HA and HA-ETMP1 strains respectively. A black arrowhead points to a specific ∼55 kDa band unique to HA-ETMP1. (C) Enriched or exclusively detected proteins in the ∼55 kDa excised gel band from HA-ETMP1 IP eluate as compared to that of mock-HA control IP eluate by LC-MS/MS sequencing analysis. MW stands for predicted molecular weight. QV denotes quantitative values (normalized total spectra). Presence of the detected proteins in the previously published mitosome proteome data (Mi-ichi et al., 2009) was performed and the result listed in the last column (+ indicates presence, - indicates absence). (D) Total cell lysates of mock-HA and ETMP1 respectively were separated by BN-PAGE, followed by anti-HA Western blot analysis. Black and red arrowheads respectively indicate the ∼180 kDa and ∼90 kDa complexes that contain HA-ETMP1.

We also performed Blue Native (BN)-PAGE analysis to assess whether ETMP1 is part of a protein complex. Anti-HA immunoblot analysis of BN-PAGE run samples indicated that HA-ETMP1 forms complexes of about 90 kDa and 180 kDa respectively (Figure 4D). Protein sequencing analysis of the excised silver-stained BN-PAGE bands containing these two complexes identified numerous proteins. Similarly, we set a cutoff value of >2.0 and the list of proteins are found in Supplementary Table S1A-C. Notably, EHD1 and its close homolog EHD3 (97% identical) were identified in both the 90 and 180 kDa complex bands. Thus, we regarded EHD1 as one of the potential interacting partners of ETMP1.

### EHD1 is an ETMP1-interacting protein that is localized to mitosomes and to vesicles of varying sizes

We expressed EHD1 in amoeba trophozoites with HA-tag at the amino terminus, as confirmed by the anti-HA immunoblot result showing a band corresponding to the expected molecular mass of HA-EHD1 (∼61 kDa) (Figure 5A). To analyze and confirm the mitosomal localization of EHD1, we performed double-staining IFA on HA-EHD1 expressing strain with anti-HA antibody and anti-APSK antiserum. We observed that the anti-HA signal is mostly localized to the membrane of vesicles of various sizes (Figure 5B). We could also notice a few punctate anti-HA signals which colocalized with the anti-APSK mitosome marker (Figure 5C, white arrowheads). Although minimal colocalization between anti-HA and anti-APSK signals were observed, some anti-APSK signals were notably seen near the vesicle membranes marked with HA-EHD1 (Figure 5C, white arrow). Immunoelectron analysis (Figure 5D) corroborate the IFA observations, as we observed mostly vesicular membrane staining of anti-HA gold particles (left panel), with occasional signals on mitosomal membranes (middle panel), some of which showed close proximity to vesicular membranes (right panel). Furthermore, immunoblot analysis of Percoll-gradient fractions indicated wide distribution of HA-EHD1 across various densities, mostly in fractions 9 to 10 and with weaker intensity in fractions 12 to 22 of the first ultracentrifugation, and in fractions A to N in the second ultracentrifugation (Figure 5E), validating the microscopic observations of HA-EHD1 vesicular and mitosomal localization.

**Figure 5.**
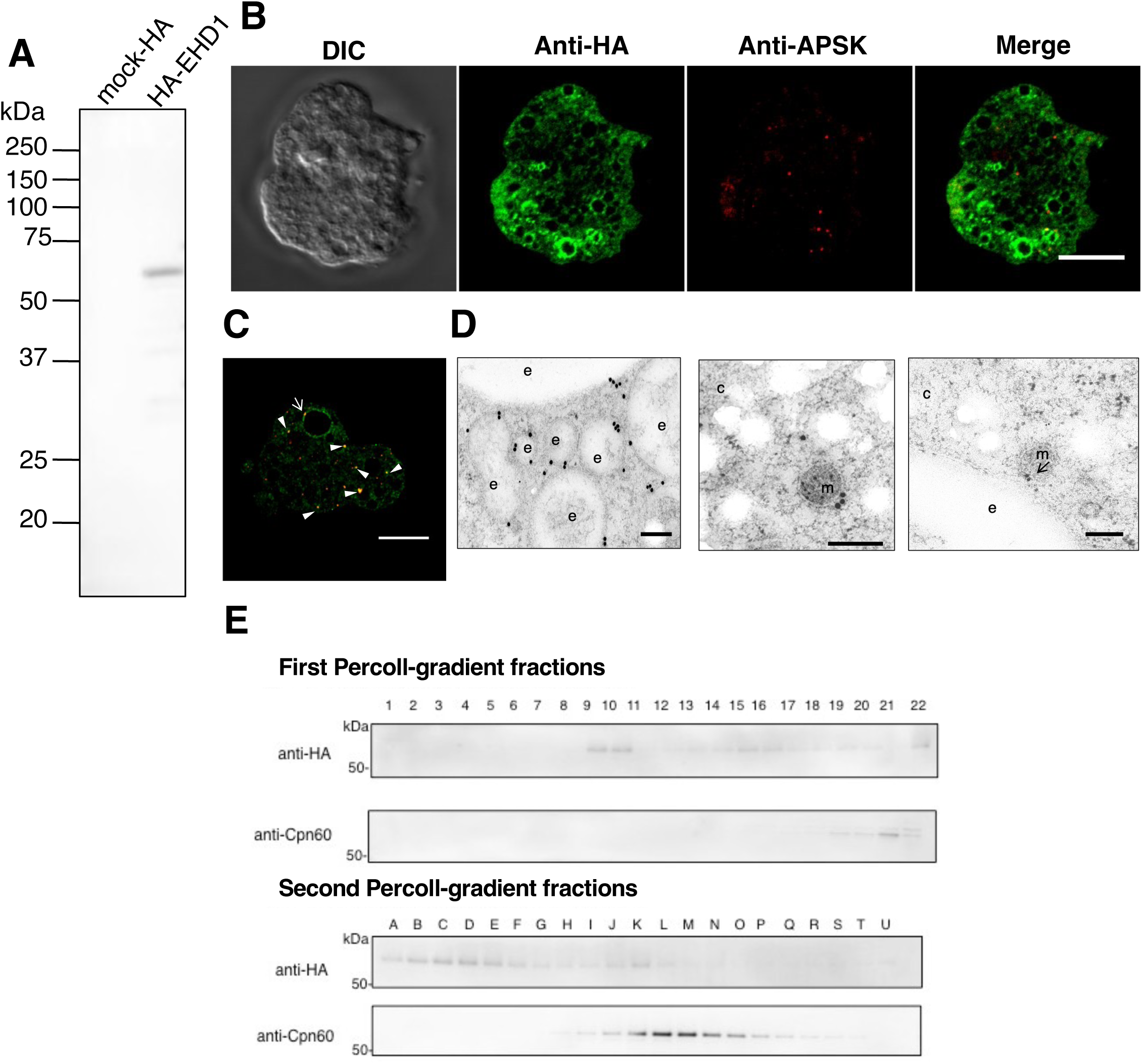
HA-EHD1 expression in *E. histolytica* trophozoites. (A) Anti-HA immunoblot analysis of approximately 30 µg total cell lysates of mock-HA and HA-EHD1, respectively, show a 61 kDa band corresponding to HA-tagged EHD1. (B-C) Representative immunofluorescence images of fixed HA-EHD1 expressing cells double-stained with anti-HA (green) and anti-APSK (red) antibodies respectively. White arrow and arrowheads indicate proximity and colocalization between anti-HA and anti-APSK signals respectively. Scale bar = 10 µm. (D) Representative immunoelectron micrographs of HA-EHD1 trophozoites, double stained with 5 nm anti-HA gold and 15 nm anti-APSK gold. Scale bar = 200 nm. The letters “c”, “e”, and “m” indicate cytosol, endosomes, and mitosomes, respectively. An arrowhead points to the structure where the membranes of the mitosome and endosome are in close contact. (E) Percoll-gradient fractionation of HA-EHD1 followed by western blot analysis using anti-HA and ant-Cpn60 antibodies respectively.

As majority of the signals of HA-EHD1 appear on vesicles, we next characterized the vesicles marked by HA-EHD1 by performing co-staining IFA using anti-HA antibody and one of the following antisera respectively: anti-vacuolar protein sorting 26 (Vps26), anti-pyridine nucleotide transhydrogenase (PNT), and anti-Rab11B. Most of the anti-HA-stained vesicles were colocalized with anti-Vps26- rather than anti-PNT- and anti-Rab11B-stained vesicles (Figure 6A) as supported by the Pearson correlation R value ranges of 0.22 to 0.37 for anti-Vps26, -0.16 to 0.19 for anti-PNT and 0.12 to 0.01 for anti-Rab11B, respectively. Vps26 is a retromer complex component and is a marker of endosomes/phagosomes in *E. histolytica* (19, 20). PNT is localized to the membrane of numerous vesicles/vacuoles, including lysosomes and phagosomes (21), while Rab11B was demonstrated to partially colocalize with late endosomes (22). Together, these data suggest EHD1 is mostly localized to endosomal membranes which may contain Vps26 and to some extent PNT, but not Rab11B.

**Figure 6.**
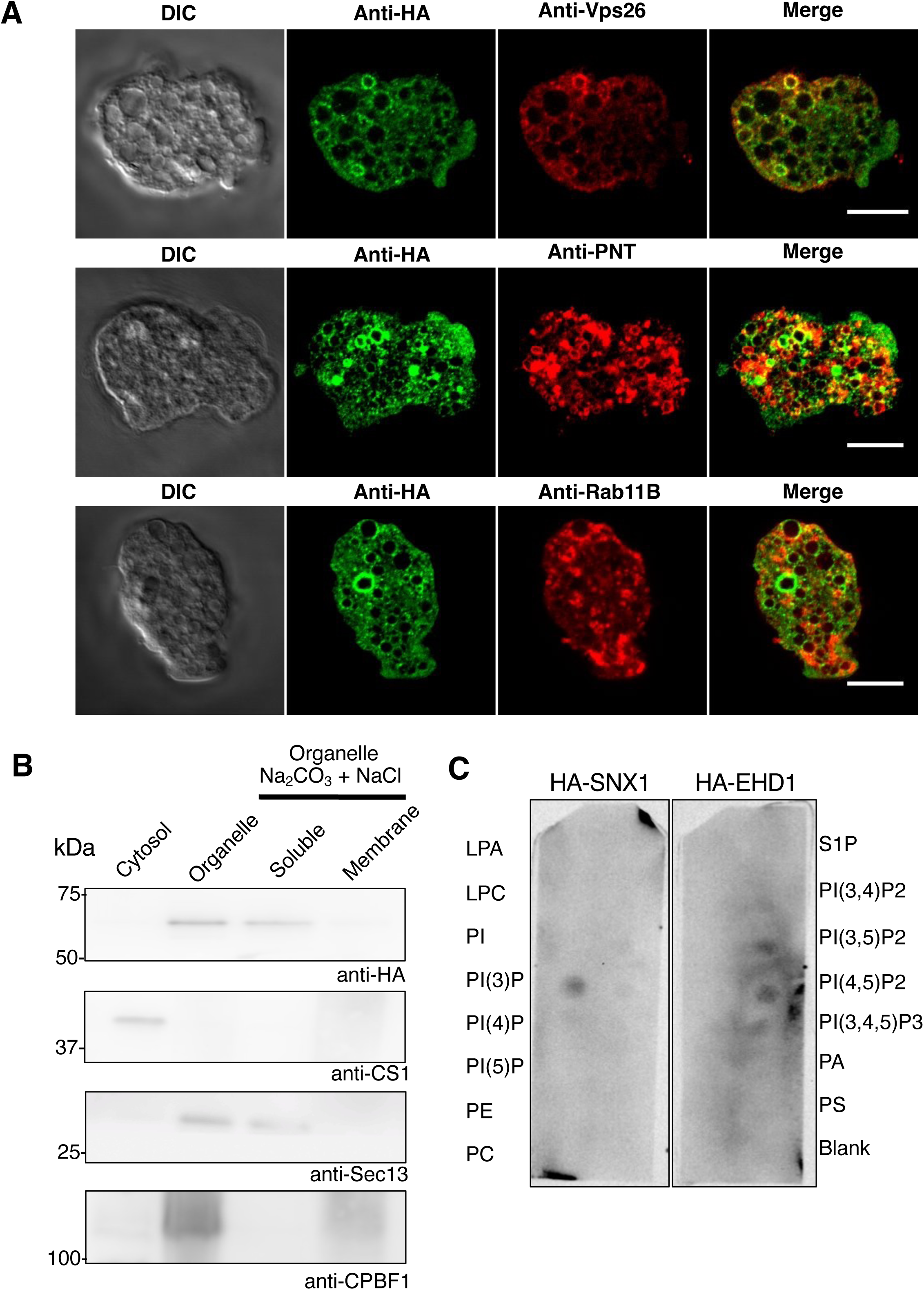
Association of HA-EHD1 to *E. histolytica* membranes. (A) Colocalization analysis of HA-EHD1 with various endosomal markers. Representative IFA images HA-EHD1 co-stained with anti-HA (green) and anti-vacuolar protein sorting 26 (Vps26, red, upper panel), anti-pyridine nucleotide transhydrogenase (PNT, red, middle panel), and anti-Rab11B (red, bottom panel) respectively. (B) Immunoblot analysis of carbonate fractionation assay of HA-EHD1 organelle-rich fraction using (from top to bottom panel) anti-HA, anti-CS1 (cytosolic protein control) anti-Sec13 (peripheral membrane protein control), and anti-CPBF1 (membrane protein control) respectively. (C) Lipid overlay assay of HA-EHD1 and HA-SNX1 (PI3P binding protein control) respectively. The membrane strips contain 100 pmol of the following lipids per spot: lysophosphatidic acid (LPA), lysophosphocholine (LPC), phosphatidylinositol (PtdIns), phosphatidylinositol (3)-phosphate (PtdIns(3)P), phosphatidylinositol (4)-phosphate (PtdIns(4)P), phosphatidylinositol (5)-phosphate (PtdIns(5)P), phosphatidylethanolamine (PE) phosphatidylcholine (PC), sphingosine 1-phosphate (S1P), phosphatidylinositol (3,4)- bisphosphate (PtdIns(3,4)P2), phosphatidylinositol (3,5)-bisphosphate (PtdIns(3,5)P2), phosphatidylinositol (4,5)-bisphosphate (PtdIns(4,5)P2), phosphatidylinositol (3,4,5)- trisphosphate (PtdIns(3,4,5)P3), phosphatidic acid (PA), phosphatidylserine (PS) respectively.

### HA-EHD1 is weakly associated to organellar membranes and preferentially binds to PI(3,5)P_2_ and PI(4,5)P_2_

We performed a similar carbonate fractionation assay to the organelle-enriched fraction of HA-EHD1 expressing strain. Based on the anti-HA immunoblots, HA-EHD1 was exclusively contained in the organelle fraction, as compared to that of the anti-CS1 profile which represents cytosolic fraction (Figure 6B). Next, we also assessed membrane integration of HA-EHD1 by carbonate treatment of the organelle-enriched fraction. Results of the immunoblots showed that HA-EHD1 is not membrane-bound as compared to the lysosomal membrane protein marker CPBF1 (Figure 6B). Instead, the profile is similar to that of the blot immunostained with an antiserum targeting Sec13, a peripheral ER membrane protein (Figure 6B). This suggests that HA-EHD1 is not organellar membrane-integrated, rather is weakly organellar membrane-associated.

To validate and characterize the phospholipid binding capacity of EHD1, we carried out a lipid overlay assay using lysates of HA-EHD1 and HA-SNX1 (PI3P binding protein control) respectively. Results indicated preferential binding of HA-EHD1 to phosphoinositide diphosphates, specifically PI(3,5)P2 and PI(4,5)P2 (Figure 6C).

### Overexpression of HA-EHD1 demonstrated enhanced multivesicular body (MVB) formation

We also expressed HA-EHD1 under the control of tetracycline (tet) induction. IFA analysis of HA-EHD1 showed the protein is similarly localized to membranes of various vesicles after 1 h and 3 h of tet induced expression respectively (Figure 7A upper and middle panel). However, at 24 h post induction with tet, we noticed drastic changes in the localization as well as in the overall intracellular vesicular patterns of expressing trophozoites (Figure 7A bottom panel), wherein large multivesicular bodies (MVBs) that were also marked with anti-HA signal were observed (Supplementary Movie S1). These findings were also supported by immunoelectron micrographs, showing immunodecoration of gold anti-HA particles along the membranes of MVBs, including the neck of invaginated vesicles (Figure 7B), after 24 h of tet-induced expression of HA-EHD1. These data point to the involvement of EHD1 in the biogenesis of MVBs in *E. histolytica*.

**Figure 7.**
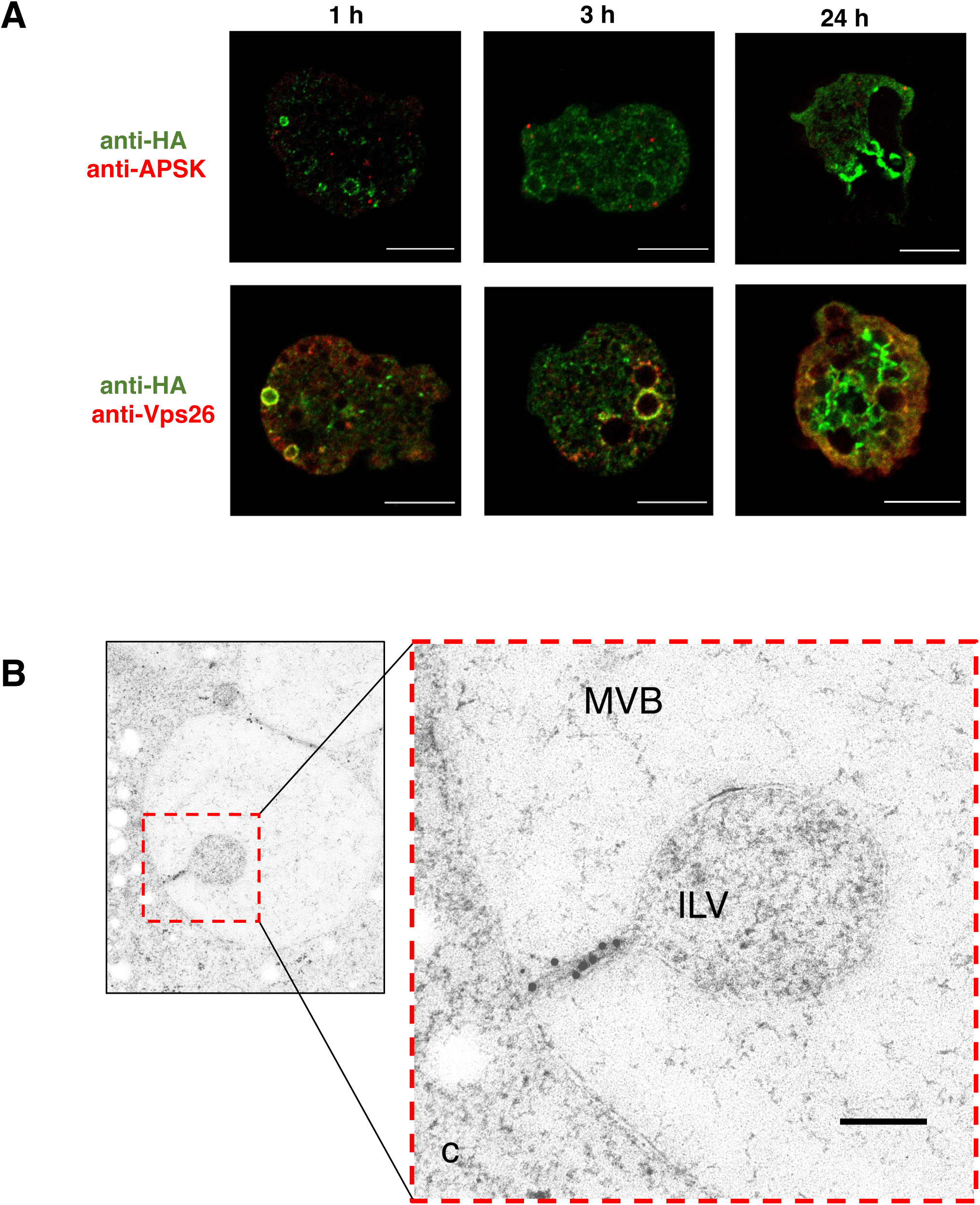
Involvement of HA-EHD1 in multivesicular body formation. (A) Representative anti-HA antibody and anti-APSK antiserum (upper panel) or anti-Vps26 antiserum (lower panel) double-staining IFA images of trophozoites that expressed HA-EHD1 trophozoites after 1h, 3h, and 24h of induction by tetracycline. Scale bar = 10 µm. (B) Representative immunoelectron image of trophozoite expressing HA-EHD1 24 h post-tetracycline induction, stained with 15 nm gold anti-HA. The initials “c”, “MVB”, and “ILV” denote cytosol, multivesicular body and intraluminal vesicle, respectively. Scale bar = 200 nm.

### EHD1 is involved in early endosome formation during macropinocytosis and receptor-mediated endocytosis

To further characterize the vesicles whose membranes are associated with EHD1, we performed endocytosis assay using either dextran conjugated to rhodamine B isothiocyanate (RITC) (for bulk endocytosis and macropinocytosis) as well as transferrin conjugated to Alexa Fluor-568 (for receptor-mediated endocytosis) respectively, as substrates, which were chased by live (GFP-EHD1 and mock-GFP) or fixed (HA-EHD1 and mock-HA) imaging analysis of treated strains. Expression of GFP-EHD1 was confirmed as a single band after anti-GFP immunoblotting (Figure 8A). From imaging of live GFP-EHD1, we observed that the GFP signals are evenly spread on round endosomal membranes (Figure 8B left panel). However, signal polarization occurs on portions where there is contact between two endosomes (Supplementary Movie S2). We also observed localization of GFP-EHD1 in endosomes that contain either RITC-dextran and Alexa Fluor 568-transferrin (Figure 8B middle and right panels respectively). Our observations also revealed that EHD1 is involved in early endosome formation during macropinocytosis of RITC-dextran. Membranes of newly formed vesicles after ingestion of RITC-dextran initially did not contain EHD1 but several seconds later, GFP-EHD1 showed intense signal on the membrane of the enclosing early endosome (Supplementary Movie S3). Consistent with this, we also noticed a similar phenomenon of GFP-EHD1 recruitment in closing early endosomes when Alexa Fluor-568-transferrin was used as substrate (Supplementary Movie S4). In addition, we observed accumulation of transferrin on to certain spots in the plasma membrane which showed remarkably high GFP-EHD1 signal (Supplementary Movie S5). This suggests that EHD1 is also involved in intra-vesicular traffic of transferrin with some aggregate signals localized near the PM, likely hinting at its involvement in receptor or membrane recycling.

**Figure 8.**
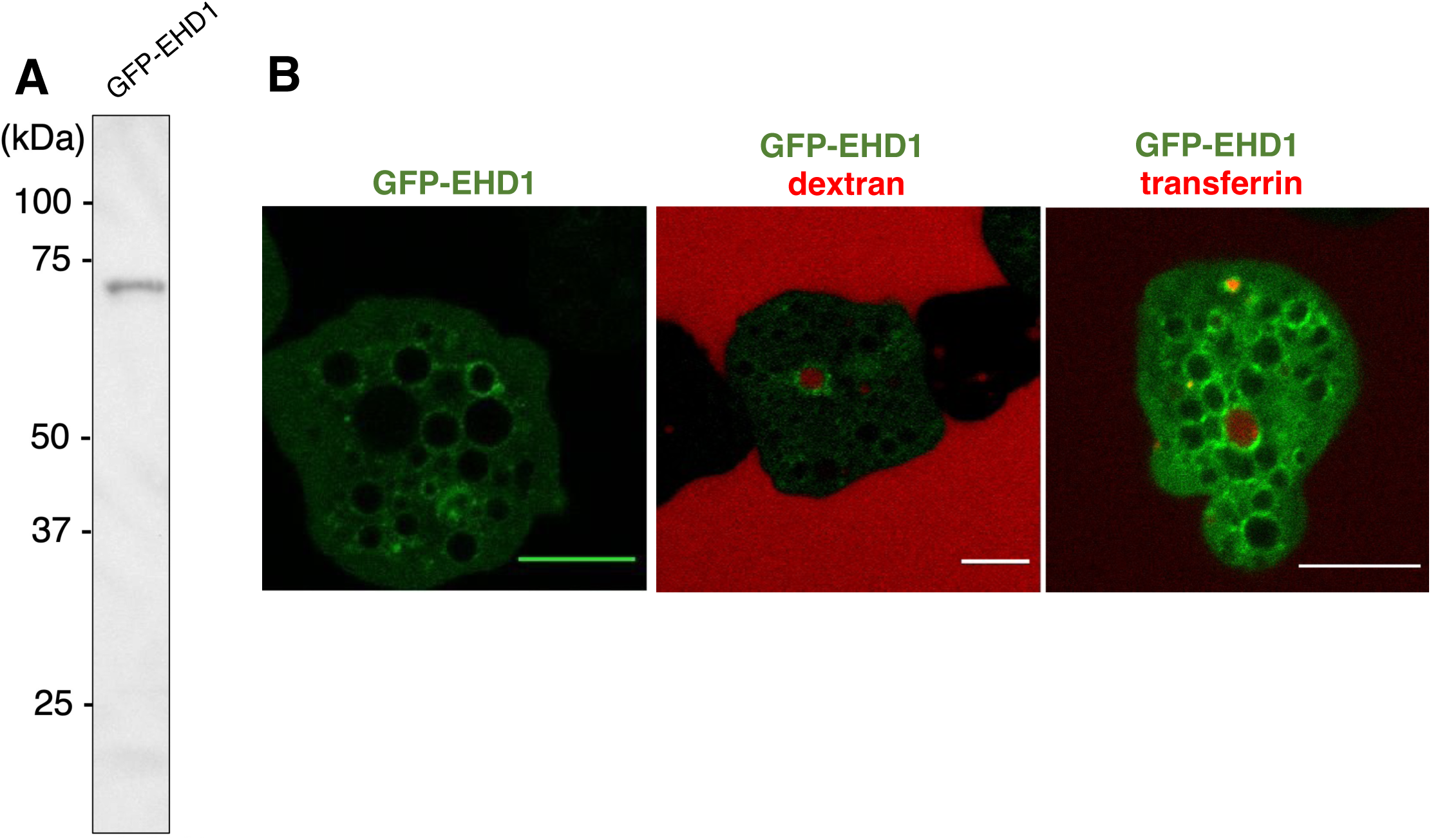
Involvement of GFP-EHD1 in amoebic endocytosis. (A) Anti-GFP immunoblot analysis of approximately 20 µg total lysate of GFP-EHD1 expressing trophozoite. (B) Confocal microscopy images from movies of live trophozoites expressing GFP-EHD1 (left panel), and GFP-EHD1 in medium supplemented with either RITC-dextran (middle panel) or Alexa Fluor 568-transferrin (right panel) respectively. Scale bar = 10 µm.

### HA-EHD1 is localized to phagosome and trogosome membrane

To further characterize EHD1-containing vesicles, we performed phagocytosis assay by co-incubating expressing trophozoites with CellTracker Blue-stained Chinese hamster ovary (CHO) cells. Live and fixed-cell imaging analyses of phagosomes or trogosomes containing whole, or bites of CHO cells, respectively were observed at varying time points after co-incubation. We observed association of either GFP-EHD1 or HA-EHD1 on some phagosome and trogosome membranes (Figure 9). We also noticed patches of higher intensity signals on certain regions of contact between phago- or trogosomes and other vesicles in both fixed (Supplementary Movie S6) and live (Supplementary Movie S7) cell imaging analyses. IFA images also suggest that HA-EHD1 is localized at the phagocytic cup/tunnel suggesting its involvement in early phagosome formation (Figure 9 top panel; 15 min post coincubation). Also observed in fixed cells was the localization of HA-EHD1 on the trogosome membrane that appears to undergo tubulation (Figure 9, bottom panel; 60 min post co-incubation).

**Figure 9.**
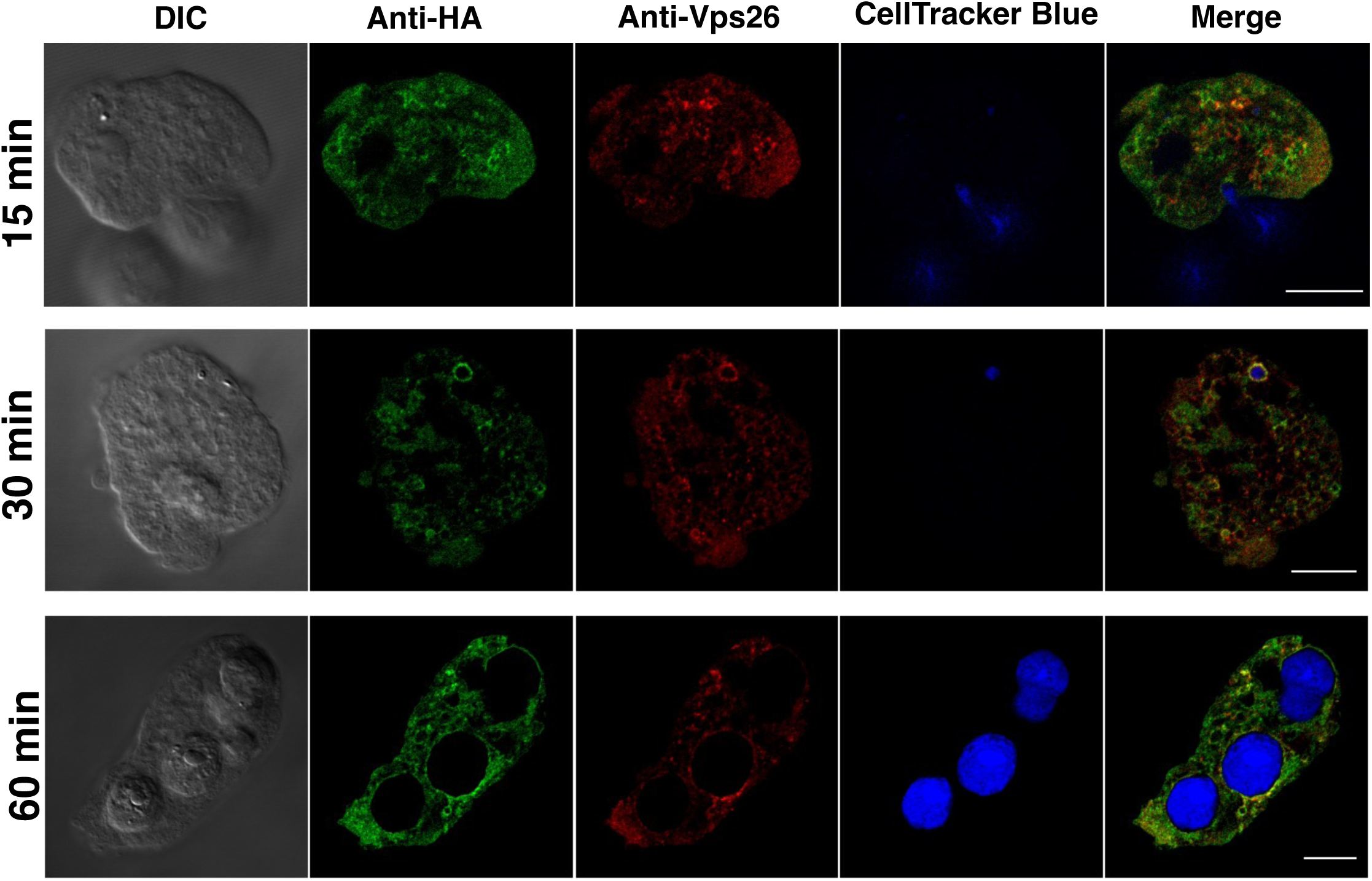
Involvement of HA-EHD1 in amoebic phagocytosis and trogocytosis. Representative IFA images of fixed anti-HA (green) and anti-Vps26 (red) double-stained HA-EHD1 trophozoites 15, 30, and 60 minutes (top to bottom) after coincubation with CellTracker Blue-stained Chinese hamster ovary (CHO) cells.

### Recombinant His-EHD1 demonstrated ATPase activity in vitro

We also expressed amino terminus histidine (His)-tagged *E. histolytica* EHD1 in bacteria to assess its enzymatic activity in vitro. We purified His-EHD1 using nickel-nitriloacetic acid (Ni-NTA)-agarose beads as shown by the Coomassie Brilliant Blue-stained SDS-PAGE gel, as well as the anti-His antibody-stained PVDF membrane (Figure 10A), containing representative Ni-NTA purification fractions. Eluted fraction of purified His-EHD1 demonstrated ATPase activity (Figure 10B) with a Michaelis-Menten constant (Km) value of 94.91 ± 16.63 μM and a maximum velocity (V_max_) of 9.85 ± 0.37 µmole/min/mg.

**Figure 10.**
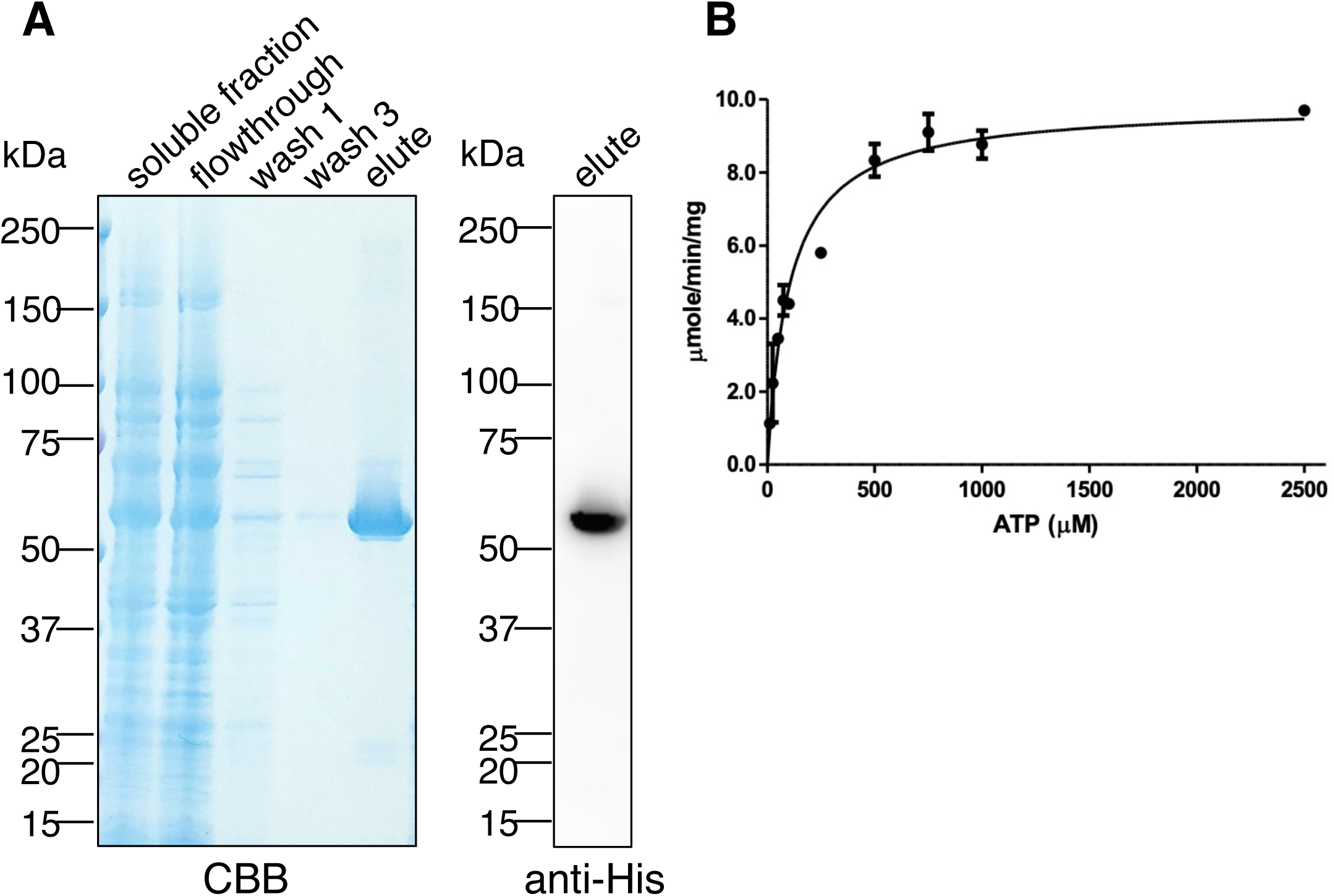
Activity assay of purified recombinant His-EHD1. (A) Coomassie Brilliant Blue-stained SDS-PAGE gel (left panel) and anti-His immunoblot (right panel) of purification fractions of His-EHD1. (C) Determination of the specific activity of His-EHD1 using ATP as substrate at various concentrations.

## Discussion

We have verified our prediction of ETMP1 being localized to the mitosomal membrane by imaging and fractionation analyses. The gene encoding for this protein is essential to the parasite’s proliferation as indicated by the failure of transfected trophozoites to survive sublethal concentration of drug pressure, compared with those transfected with an empty vector control. Previous attempts at silencing the genes encoding other mitosomal membrane proteins such as Tom40 (23) and MBOMP30 (24) also failed, suggesting the essential role that these proteins, and the mitosome itself where they exclusively localize, maintain in the proliferation of *E. histolytica*. Overexpression of ETMP1 also affected the growth rate of the parasite negatively. This may be due to the disruption of tight regulatory mechanisms for maintaining mitosomal homeostasis and/or formation of toxic protein aggregates. It could also be due to the stoichiometric imbalance of HA-ETMP1-containing protein complexes. Our BN-PAGE analysis identified ETMP1 in the 90kDa and 180 kDa complexes respectively, whose formation, compositional ratios, and biological functions may be sensitive to ETMP1 overexpression.

HA-ETMP1 immunoprecipitated a unique ∼55 kDa protein. Mass spectrometry analysis of the excised silver-stained gel band indicated several candidates including EH-domain containing protein (EHD1; EHI_105270; 58 kDa) and its ortholog (EHD2; EHI_152680; 58 kDa) sharing 82% identity, vacuolar protein sorting-associated protein 45 (60 kDa; EHI_154290), and L-myo-inositol-1-phosphate synthase (57 kDa; EHI_070720). Incidentally, when we sequenced the complex band of 90 and 180 kDa BN-PAGE complex bands that include HA-ETMP1, we identified EHD1 and its close homolog EHD3 (EHI_052870; 58 kDa) with 97% identity. From these data, we deduced a plausible interaction between ETMP1 and EH domain containing proteins, with a focus on EHD1 in this paper. Repeated multiple attempts at immunoprecipitating the said complexes failed. One possibility is that the topology of HA-ETMP1 in the complex blocked the HA epitope tag from binding to the anti-HA beads. We also performed IP using HA-EHD1 (Supplementary Figure S1, Supplementary Table S2A), however our protein sequencing analysis of the ∼30-37 kDa excised band did not detect HA-ETMP1 (Supplementary Table S2B), suggesting the likely transient nature of this protein binding. The detection of amoebic EHD isotypes in the pull-down and BN-PAGE complexes of HA-ETMP1 respectively, suggests potential interaction among these EHD homologs. It is also plausible that amoebic EHDs form hetero-dimers or hetero-oligomers as was demonstrated by mouse EHD1 and EHD3. The interaction between mouse EHD1 and EHD3 is likely involved in the regulation of recycling endosomes movement along microtubules (25). In *E. histolytica,* such EHD oligomers may be involved not only during endocytosis but may also exist during the formation and maintenance of the mitosome-endosome contact. Compositional variations of EHD homo- or hetero-oligomers may also exist, and their corresponding functions may be stoichiometry-dependent.

EHDs have been associated to play roles in various endocytic processes. In one subset known as the C-terminal EHDs, four paralogues are present in mammals, namely EHD1, EHD2, EHD3, and EHD4. Mammalian EHD1 regulates exit of proteins from the endocytic recycling compartment to the plasma membrane, while both EHD1 and EHD3 have similar roles in controlling early endosome to Golgi transport (26, 27). Mammalian EHD2 localizes to caveolae and together with the Bin Amphiphysin Rvs (BAR)-domain containing binding partner PACSIN2, stabilizes caveolae at the cell surface (28), whereas mammalian EHD4 facilitates macroendocytic uptake of tropomyosin receptor kinase (Trk) receptors (29). EHDs are also implicated in the regulation of endocytic pathways associated with lipid metabolism. Mammalian EHD1 is involved in cholesterol homeostasis, affecting generation of cholesterol and triglyceride lipid bodies (30).

EHDs also regulate endocytosis in other organisms including plants, worms, and protozoans. *Arabidopsis thaliana* has two EHD paralogs, *At*EHD1 and *At*EHD2. Downregulation of *At*EHD1 led to a deficiency in the entry of endocytosed material into plant cells, whereas overexpression of *At*EHD2 caused an inhibitory effect on endocytosis, suggesting both proteins are important components in plant endocytic machinery (31). The EHD ortholog in *Caenorhabditis elegans*, receptor-mediated endocytosis 1 (Rme1), localizes to the endocytic recycling compartment and mediates the exit of cargo proteins to the cell membrane (32). In the protozoan parasite that causes malaria, *Plasmodium falciparum,* a single EHD protein is encoded in its genome. *Pf*EHD is involved in endocytosis and plays a role in the generation of endocytic vesicles at the plasma membrane, that are subsequently targeted to the neutral lipid generation/storage site localized near the food vacuole (33). In the free-living amoebozoan *Dictyostelium discoideum*, a single gene encoding EHD protein was identified. *Dd*EHD was determined to be involved in phagosome maturation, and its deletion resulted to defects in intraphagosomal proteolysis and acidification, early delivery of lysosomal enzymes, and fast retrieval of the vacuolar H^+^-ATPase in maturing phagosomes (34).

We have shown that *Eh*EHD1 is involved in various endocytic processes. Our live imaging analysis showed involvement of *Eh*EHD1 in early endosome formation, particularly during closure of newly-formed endosomes after engulfment of either RITC-dextran (Supplementary Movie S3) or Alex568-transferrin (Supplementary Movie S4), suggesting that *Eh*EHD1 may participate in the scission of early endosomes generated from macropinocytosis as well as receptor-mediated endocytosis. Vesicle tubulation and scission are associated roles of EHDs as they possess a dynamin-like ATPase domain as demonstrated previously in (34–37).

Our in vitro enzyme assay showed that His-EHD1 has ATPase activity with a Km value of 94.91 ± 16.63 μM as compared to the previously reported Km values for mouse EHD1 (80 μM) and *Ce*RME1 (30-μM) (38). We also attempted to investigate the role of ATPase activity of EHD1 in *E. histolytica* by expressing ATPase-deficient dominant negative mutant, however the transfectants did not survive drug selection, suggesting the importance of EHD1 ATP hydrolysis in amoebic biology. Based on other works, ATPase activity of C-terminal EHD-containing proteins is crucial for various stages of the endocytic traffic machinery. Hydrolysis of ATP was essential for binding of human EHD2 complexes to caveolae during clathrin-independent endocytosis (39). It is suggested that membrane scission results from ATP hydrolysis by human EHD2 in vivo (35). Using cross-complementation assays in *C. elegans*, ATP binding and hydrolysis of human EHD1 was essential for endocytic recycling. It was also shown using in vitro liposome-based assays that ATP binding of human EHD1 promote scaffold self-assembly, while ATP hydrolysis enables extension of bulges and thinning of tubular model membranes which leads to scission (40). In vitro analysis also revealed that ATP binding and concomitant hydrolysis allows membrane remodelling into highly curved tubules (29). We can only hypothesize that ATP hydrolysis in amoebic EHD1 may have functions similar to its homologs in other organisms.

We also detected *Eh*EHD1 in the phagocytic cup, and membranes of phagosomes and trogosomes, although only a select few phagosomes and trogosomes are labeled with EhEHD1 in both live and fixed imaging analysis. The same can also be said when we performed endocytosis assay using either RITC-dextran or Alexa Fluor 568-transferrin. This suggests the nature of EHD localization being dependent on either recruitment by interacting proteins or association/binding with certain lipids on vesicular membranes at specific time points. This is reflected by the localization of either GFP-EHD1 or HA-EHD1 in membranes of vesicles of various sizes, and the seemingly polarized signal intensity onto sites where two vesicles are in close contact.

As suggested by our lipid overlay assay result, *Eh*EHD1 preferentially binds to PI(3,5)P2 and PI(4,5)P2. PI(4,5)P_2_ has been demonstrated to be localized to the plasma membrane (41), lipid rafts and uroids (42) of *E. histolytica.* It is important to note that PI(4,5)P2 localized at the plasma membrane is involved initiating internalization during endocytosis, micropinocytosis, and phagocytosis (43, 44), whereas PI(3,5)P2 has a critical role in endosome/lysosome biogenesis, and in the initiation of MVB formation (45). Together, these results circumstantially support our observations of amoebic EHD1 localization and involvement in early endosome, intraluminal vesicle, and MVB formation.

On the possible role(s) of mitosome-endosome contact in *E. histolytica*, we posit that this MCS may be involved in lipid transfer, ion transport, and quality control. Lipid transport and/or metabolism are commonly alluded roles of MCSs. Although we did not detect any lipid transport proteins in our immunoprecipitation assay, two lipid transport proteins (LTP1 and LTP3) in *E. histolytica* have been characterized (46), and it is plausible that various LTPs may transiently interact with amoebic MCSs to facilitate lipid mobility across organelles. We detected a few fatty acid ligases in the ∼90 and ∼180 kDa complex, however the interaction of these proteins to the HA-ETMP1 containing complex needs to be experimentally validated. Alternatively, ion transport may also be facilitated in this MCS, as was demonstrated in epithelial cells, where the mitochondria and endosomes that contain iron-bound transferrin are involved in “kiss and run” interactions, leading to iron transfer from endosomes to mitochondria (47). Another possibility is the involvement of EHD1 in mitosomal dynamics. Mitochondria undergo dynamics of fusion and fission to ensure maintenance of homeostasis, control of reactive oxygen species, apoptosis, and autophagy. Dynamin and dynamin related proteins (Drps) have been implicated in mitochondrial fission. Recently in HeLa cells, EHD1 was reported to be a novel regulator of mitochondrial fission via a mechanism distinct from that of dynamin/Drp. In this model human EHD1, together with its binding partner rabankyrin-5 interact with the retromer complex participate in mitochondrial division. EHD1 was suggested to facilitate the fission of vesicles that transport Vps35, a retromer complex component, from endosomes to the mitochondrial membrane. It was also suggested that Vps35 may interact with inactive Drp1 on the mitochondrial membrane, causing its removal and subsequent action of active Drp1 to perform mitochondrial fission (48). Fission has also been reported in MROs of anaerobic parasites such as the hydrogenosomes of *Trichomonas vaginalis* and *E. histolytica* (49–51). Mitosome fission in *E. histolytica* was reported to involve a heterodimer complex of two dynamin-related proteins, DrpA and DrpB (50). It is interesting if amoebic EHD1 also takes part in influencing mitosome fission as was postulated in mammalian cells (48). An alternative novel pathway for mitochondrial quality control that is independent of Atg5 and LC3 is the formation of mitochondria-derived vesicles targeted to lysosomes. Ultrastructural analysis of COS7 cells identified the presence vesicles that are Tom20-positive within MVBs (52). Furthermore, in hepatocytes, a complex made up of EHD2, EH domain binding protein 1 (EHBP1), and Rab10, promotes extension of the LC3-containing autophagic membrane in order to engulf lipid droplets during lipophagy (53). Such related pathways may also exist in *E. histolytica* that warrants further investigation in the future.

## Conclusion

We report a novel membrane contact site between mitosomes and endosomes of *Entamoeba histolytica*. This unprecedented MCS features the mitosomal membrane protein ETMP1 and a C-terminal EH domain containing protein, EHD1. ETMP1 is a protein unique to *Entamoeba* and is essential to parasite proliferation. It interacts with EHD1, a protein involved in various endocytic processes in *E. histolytica*, namely in early endosome formation during bulk and receptor-mediated endocytosis, phagocytosis and trogocytosis of mammalian cells, and in the invagination of intraluminal vesicles for the generation of multivesicular bodies. Such novel ETMP1-EHD1 interaction hints at a possible role of this mitosome-endosome MCS on various physiological processes that have been demonstrated in other organisms. We thus propose that ETMP1-EHD1 mediated contact site is involved in lipid transfer, biogenesis, autophagy, organelle dynamics and quality control of MROs. Further investigation is needed to fully dissect the molecular mechanisms and functions of this and other MRO-related MCSs.

## Materials and Methods

### Entamoeba histolytica cultivation

*Entamoeba histolytica* HM-1:IMSS Cl6 (54) and G3 (55) strains were maintained in Diamond’s BI-S-33 medium (54) as described previously. Subculture was performed after incubation of up to 3-4 days when trophozoites reached the late-logarithmic phase.

### Plasmid construction

Extraction of total RNA from *E. histolytica* trophozoites, purification of mRNA, and synthesis of cDNA were performed by following protocols described previously (24). For the expression of hemagglutinin (HA) tagged proteins in *E. histolytica* trophozoites, target genes (*etmp1*: EHI_175060 and *ehd1*: EHI_105270) were amplified by polymerase chain reaction (PCR) using *E. histolytica* cDNA as template, and the corresponding primer sets: (etmp1-*Xma*I-fwd: GTTcccgggATGGAACAAATAACTGAAGAA; *etmp1*-*Xho*I-rev: GAActcgagTTATTTTTTCATTTTTCTTAAGG; and *ehd1*-*Xma*I-fwd: GTTcccgggATGTTTGGTAAGAAGAAACAAAAACC; *ehd1*-*Xho*I-rev: GAActcgagTTATTCAACTGGTGGAAGATTGTC). These PCR amplicons were inserted into the following plasmids: pEhEx-HA and pEhEx-GFP for constitutive expression, (56) and pEhtEx-HA and pEhtEx-GFP for tetracycline-induced expression (50), after digestion with *Xma*I and *Xho*I (New England Biolabs, Beverly, MA, USA) and then ligated using Ligation Convenience Kit (Nippongene, Tokyo, Japan). For the expression of recombinant proteins in *Escherichia coli*, PCR-amplification of *ehd1* was performed using *E. histolytica* cDNA as template and the following primer set (*ehd1*-*Bam*HI-fwd: GTTggatccATGTTTGGTAAGAAGAAACAAAAACC, and *ehd1*-*Sal*I-rev: GAAgtcgacTTATTCAACTGGTGGAAGATTGTC). Digestion and ligation to *Bam*HI and *Sal*I-linearized pColdI plasmid (Takara, Shiga, Japan) were performed. For transcriptional gene silencing, about 400 bp fragments of *etmp1* and *ehd1* were amplified using cDNA and appropriate primer sets: etmp1gs-*Stu*I-fwd: GTTaggcttATGGAACAAATAACTGAAG; etmp1gs-*Sac*I-rev: GAAgagctcCTAATTTGATTCCTTTTAAAG; and ehd1gs-*Stu*I-fwd: GTTaggcctATGTTTGGTAAGAAGAAACAA and ehd1gs-*Sac*I-rev: GAAgagctcTAAATTTAGCCATAAATTCAT. The amplicons were digested with *Stu*I and *Sac*I and ligated to pSAP2-Gunma (57).

### Amoeba transfection and drug selection

The constructed plasmids described above were transfected by lipofection into *E. histolytica* trophozoites, as described previously (58, 59). Selection of transfectants was performed by changing the culture medium supplemented with G418 (Gibco/Life Technologies, USA) for those transfected with pEhEx-based plasmids, or with hygromycin (Fujifilm Wako, Japan) for those transfected with pEhtEx-based plasmids. The starting concentration of either 1 µg/mL G418 or hygromycin added was gradually increased until all control cells (transfected without plasmid) died from the antibiotic challenge. All resultant strains were maintained in medium containing 10 µg/mL G418 or 20 µg/mL hygromycin unless otherwise stated. For tetracycline-induction of protein expression, 10 µg/mL tetracycline was added to semi-confluent cultures 24 h prior to performing assays unless otherwise stated.

### Immunoflourescence assay (IFA)

Double-staining immunofluorescence assay was performed as previously described (13), using anti-HA mouse monoclonal antibody (clone 11MO, Covance, USA) diluted 1:500 in 2% saponin and 0.1% bovine serum albumin in phosphate buffered saline (saponin-BSA-PBS), to detect HA-tagged ETMP1 and EHD1, respectively, and one of the following polyclonal rabbit antisera diluted 1:1000 in saponin-BSA-PBS unless otherwise stated: anti-adenosine-5’-phosphosulfate kinase (APSK; EHI_179080; a mitosomal matrix protein; (57) diluted 1:300; anti-vacuolar protein sorting 26 (Vps26; EHI_062490; a retromer complex component (19)); anti-Rab11B (EHI_107250; involved in cysteine protease secretion; (22)); and anti-pyridine nucleotide transhydrogenase (PNT; EHI_014030; a novel class of lysosomal PNT; (21)). Secondary antibodies used were Alexa Fluor-488 anti-mouse antibody and Alexa Fluor-568 anti-rabbit antibody Thermo Fisher diluted 1:1000 in saponin-BSA-PBS, respectively. Cells were visualized using LSM780 (Carl Zeiss Microscopy, Germany) confocal laser scanning microscope.

### Subcellular fractionation and immunoblot analysis

Trophozoites at the late-logarithmic phase were collected and washed thrice with 2% glucose-PBS. Cells were mechanically disrupted using a Dounce homogenizer as described previously (13). The resulting homogenate was separated by Percoll-gradient fractionation as previously described (13, 18). For carbonate fractionation, organelle-enriched fractions from HA-ETMP1, MBOMP30-HA, HA-EHD1 and mock control homogenates were collected by centrifugation at 100,000 *g* for 60 min at 4°C. The resultant pellet was reacted with sodium carbonate as previously described (13, 23, 24). All fractions collected were run in SDS-PAGE followed by Western blotting as previously described (60). Immunostaining of PVDF membranes was performed using anti-HA antibody, anti-APSK antiserum (organelle fraction marker), anti-cysteine synthase 1 (CS1; EHI_171750; cytosolic enzyme involved in cysteine metabolism) (61), anti-cysteine protease binding family protein 1 (CPBF1; EHI_164800, membrane fraction control) (62) and chemiluminescent bands were visualized using LAS-4000 mini luminescent image analyzer (Fujifilm Life Science, Tokyo, Japan).

### In silico predictions and analyses

Transmembrane domain-containing mitosomal proteins were predicted using a pipeline developed in our previous study (12). To search for homologs of ETMP1 in various *Entamoeba* species, we used as query the *E. histolytica* protein, EHI_175060, and implemented a BLAST search using the *Amoebozoa* resource database, AmoebaDB (63). Coiled-coil regions were predicted using DeepCoil (64).

### Immunoelectron microscopy

Samples were prepared as described previously (24). The specimens were double-stained with anti-HA mouse antibody, and anti-APSK rabbit antiserum (57). Processing and visualization were performed by Tokai Microscopy Inc. (Nagoya, Japan), using a transmission electron microscope (JEM-1400 Plus, JEOL Ltd., Japan) at an acceleration voltage of 80 kV. Digital images with a resolution of 2048 × 2048 pixels were taken using a CCD camera (VELETA, Olympus Soft Imaging Solution GmbH, Germany).

### Immunoprecipitation (IP) of HA-ETMP1 by anti-HA antibody

Organelle-enriched fractions from HA-ETMP1 and mock pEhEx-HA control homogenates were prepared and approximately 2 μg of proteins were solubilized in 2% digitonin in IP Buffer containing 50 mM BisTris-HCl, pH 7.2, 50 mM NaCl, 0.001 % Ponceau S, and 10 % w/v glycerol for 30 min on ice. The solubilized fraction was collected by centrifugation at 20,000 *g* for 30 min at 4 °C. Immunoprecipitation was performed as previously described (23). Bound proteins were eluted overnight using 60 μg HA peptide. Eluted fractions were loaded on SDS-PAGE gels, followed by immunoblotting using mouse anti-HA antibody. Silver staining was performed using the Silver Stain MS kit (Fujifilm Wako Pure Chemical Corporation, Osaka, Japan), according to the manufacturer’s protocol. Protein sequencing by liquid chromatography-mass spectrometry analysis was conducted by the Biomolecular Analysis Facility Core, University of Virginia.

### Lipid overlay assay

As described previously (20), the lysate of HA-EHD1 expressing strain was used to probe a P-6001 phospholipid membrane strip (Echelon Biosciences, Salt Lake City, Utah, USA). The lysate of HA-SNX1 which binds to PI3P (20) was used as a positive control. The strips were washed three times with 0.1% Tween 20 in PBS (PBS-T), followed by reaction with 1:1000 anti-HA mouse antibody in 3% BSA-PBS for 2 h at room temperature. The strips were washed and incubated with 1:6000 HRP-conjugated goat anti-mouse IgG (Thermo-Fisher Scientific, USA) in 3% BSA-PBS for 1 h at room temperature. Finally, the strips were washed and reacted with the Immobilon ECL Ultra Western HRP Substrate (Millipore, USA) following manufacturer’s instructions.

### Endocytosis assay

Approximately 1×10^5^ GFP-EHD1 or mock-GFP expressing trophozoites in 1 mL BI-S-33 of strains respectively were placed on a 35 mm collagen-coated glass-bottom culture dish (MatTek Corporation, Ashland, MA) for 15 minutes to allow for cell attachment. The medium was removed and replaced with 1 mL of BI-S-33 supplemented with either 2 mg/mL RITC-dextran (MW = 70 000; Sigma-Aldrich, USA) or 100 μg/ml Alexa Fluor-568 transferrin (Thermo-Fisher Scientific, USA). Chase was performed for up to 30 min for live imaging. For IFA, fixation was conducted onto HA-EHD1 and mock-HA strains after 0, 30, 60, 120 min of addition of either RITC-dextran or Alexa Fluor 568-transferrin. Images were captured using a confocal laser scanning microscope LSM780 (Carl Zeiss Microscopy, Germany), as the cells were being incubated at 35 °C using a temperature-controlled stage plate (Carl Zeiss Microscopy, Germany).

### Phagocytosis assay

A semi-confluent culture of Chinese hamster ovary (CHO) cells, grown in F12 medium (Sigma-Aldrich, USA) was stained by addition of 40 μM CellTracker Blue (Thermo-Fisher Scientific, USA) for 30 min at 37°C. The medium containing excess dye was removed and the cells were washed in 1X PBS followed by treatment with 0.1% trypsin for 5 min at 37°C. The detached cells were collected and washed with 1X PBS thrice by centrifugation at 3000 rpm for 3 minutes. Stained CHO cells were resuspended in BI-S-33 medium prior to addition to amoeba cells. Cells were co-incubated for 15, 30, and 60 min respectively, after which they were fixed for IFA analysis as mentioned above, but using anti-GFP antibody (Sigma-Aldrich, USA) and anti-Vps26 antiserum respectively. A parallel setup was prepared for live imaging analysis.

### Expression and purification of recombinant His-EHD1

*Escherichia coli* BL21 strain was transformed using the pCold-His-EHD1 plasmid described above, and the transformants were selected using LB agar containing 150 μg/ml of ampicillin. Isolated colonies were cultured in LB medium with 150 μg/ml of ampicillin, incubated at 37 °C with shaking. A 1L culture was inoculated and incubated in a shaker at 37 °C until reaching optical density (OD) 600 of 0.7. The culture was flash cooled in an ice water bath for 30 minutes. Induction of protein expression was made by adding 0.5 mM isopropyl β-D-thiogalactopyranoside (IPTG) to the medium followed by incubation at 15 °C with shaking for 24 h. Cells were collected and protein expression was confirmed by loading the soluble and insoluble fractions in SDS-PAGE followed by Coomassie blue staining and anti-His immunoblot analysis respectively as described previously (65). His-EHD1 was purified by binding with Ni^2+^-NTA His-bind slurry (Qiagen, Germany) and eluting with imidazole as described previously (65). Purified His-EHD1 was stored at -80°C with 20% glycerol in small aliquots until use.

### Enzyme activity assay

Varying amounts of purified His-EHD1 (0, 0.125, 0.25, 0.5, 1.0 μg) were resuspended in assay buffer (20 mM HEPES pH 7.5, 0.005 % Tween 20, 10 % glycerol, 1 mM DTT, 20 mM NaCl, and 10 mM MgCl_2_) and loaded triplicate onto independent wells of a 96-well plate. Then, 2 μl of 100 mM ATP was used as substrate and distilled water was added to bring the volume of the mixture to 20 μl. Finally, 20 μl of 2X stock solution (66) containing 100 mM Tris-HCl (pH 7.5), 10 mM MgCl_2_, 0.02% Triton X-100, 0.01% BSA, 2 mM glucose, 0.2 mM NADP, 2 u/mL ADP-hexokinase, 2 U/mL glucose-6-phosphate dehydrogenase, 2 U/mL diaphorase I, 0.1 mM resazurin in DMSO, and 20 mM N-ethylmaleimide in DMSO, was added and the plate was incubated at 37 °C for 30 min. For determining kinetic parameters, 0, 12.5, 25, 50, 75, 100, 125, 250, 500, 750, 1000, and 2500 μM ATP was used to react with 1.5 μg His-EHD1 for 30 min. The fluorescence was measured continuously at excitation and emission wavelengths of 540 nm and 590nm, respectively using SpectraMax Paradigm Multi-Mode microplate reader (Molecular Devices, San Jose, CA, USA).

## Acknowledgments

This research is funded by Grants-in-Aid for Scientific Research (B) (JP18H02650 and JP21H02723 to T.N.), Grants-in-Aid for Young Scientists (JP20K16233 to H.J.S.) and Core-to-Core Program, (JPJSCCB20190010) from the Japan Society for the Promotion of Science, Grant for research on emerging and re-emerging infectious diseases from Japan Agency for Medical Research and Development (AMED, JP20fk0108138 to T.N.). The authors want to thank Mihoko Imada of The University of Tokyo for her techincal support. The authors also want to thank Dr. Takashi Makiuchi of Tokai University School of Medicine, Japan, and Dr. Eri Hayakawa of Jichi Medical University, Japan for their valuable discussion.

## Supplemental Materials

**Supplementary Movie S1**

Multiple z-section images of fixed HA-EHD1, 24 h after tetracycline induction. Green and red signals indicate anti-HA and anti-APSK antibodies respectively. Scale bar = 5 µm.

**Supplementary Movie S2**

Live imaging of GFP-EHD1 after 24 of tetracycline induction. Scale bar = 5 µm.

**Supplementary Movie S3**

Live imaging of GFP-EHD1 trophozoites chased a few minutes after addition of RITC-dextran. Note the recruitment of GFP-EHD1 in newly closed endosomes. Scale bar = 5 µm.

**Supplementary Movie S4**

Live imaging of GFP-EHD1 trophozoites chased a few minutes after addition of Alexa Fluor 568-transferrin. Note the recruitment of GFP-EHD1 in newly closed endosomes.

**Supplementary Movie S5**

Live imaging of GFP-EHD1 trophozoites chased a few minutes after addition of Alexa Fluor 568-transferrin. Note the accumulation of GFP-EHD1 in the plasma membrane where aggregated transferrin is located. Scale bar = 5 µm.

**Supplementary Movie S6**

Multiple z-section images of fixed GFP-EHD1, (60 min after co-incubation with CellTracker blue-stained Chinese hamster ovary cells. Scale bar = 5 µm.

**Supplementary Movie S7**

Live imaging of GFP-EHD1 trophozoites chased a few minutes after co-incubation with CellTracker blue-stained Chinese hamster ovary cells. Scale bar = 5 µm.

**Supplementary Figure S1**

**Anti-HA immunoprecipitation of HA-EHD1 and mock-HA control.** (A) Western blot analysis of various IP fractions probed using anti-HA antibody (B) Silver-stained SDS-PAGE gel showing separated protein bands from eluted IP samples. Black boxes indicated regions excised and submitted for subsequent protein sequencing analysis.

**Supplementary Table S1**

(A) Exclusively detected proteins in the ∼180 kDa excised blue-native PAGE gel band from HA-ETMP1 as compared to that of mock-HA control by LC-MS/MS sequencing analysis. MW indicates predicted molecular weight, while QV denotes quantitative values (normalized total spectra). (B) Detected proteins in the ∼180 kDa excised blue-native PAGE gel band enriched in HA-ETMP1 as compared to that of mock-HA control by LC-MS/MS sequencing analysis (Qv HA/ETMp1/mock-HA ≥ 2.0.) (C) Exclusively detected proteins in the ∼90 kDa excised blue-native PAGE gel band enriched in HA-ETMP1 as compared to that of mock-HA control by LC-MS/MS sequencing analysis.

**Supplementary Table S2**

(A) Proteins identified in the 55-58 kDa excised gel band of HA-EHD1 and mock-HA IP eluate samples respectively, by LC-MS/MS analysis. MW indicates predicted molecular weight, while QV denotes quantitative values (normalized total spectra). (B) Proteins identified in the 30-33 kDa excised gel band of HA-EHD1 and mock-HA IP respectively, by LC-MS/MS analysis. MW indicates predicted molecular weight, while QV denotes quantitative values (normalized total spectra).

