## Supplementary figures and images for "*Entamoeba histolytica* EHD1 is involved in mitosome-endosome contact"

### Supplementary Figure S1

#### Supplementary Figure S1

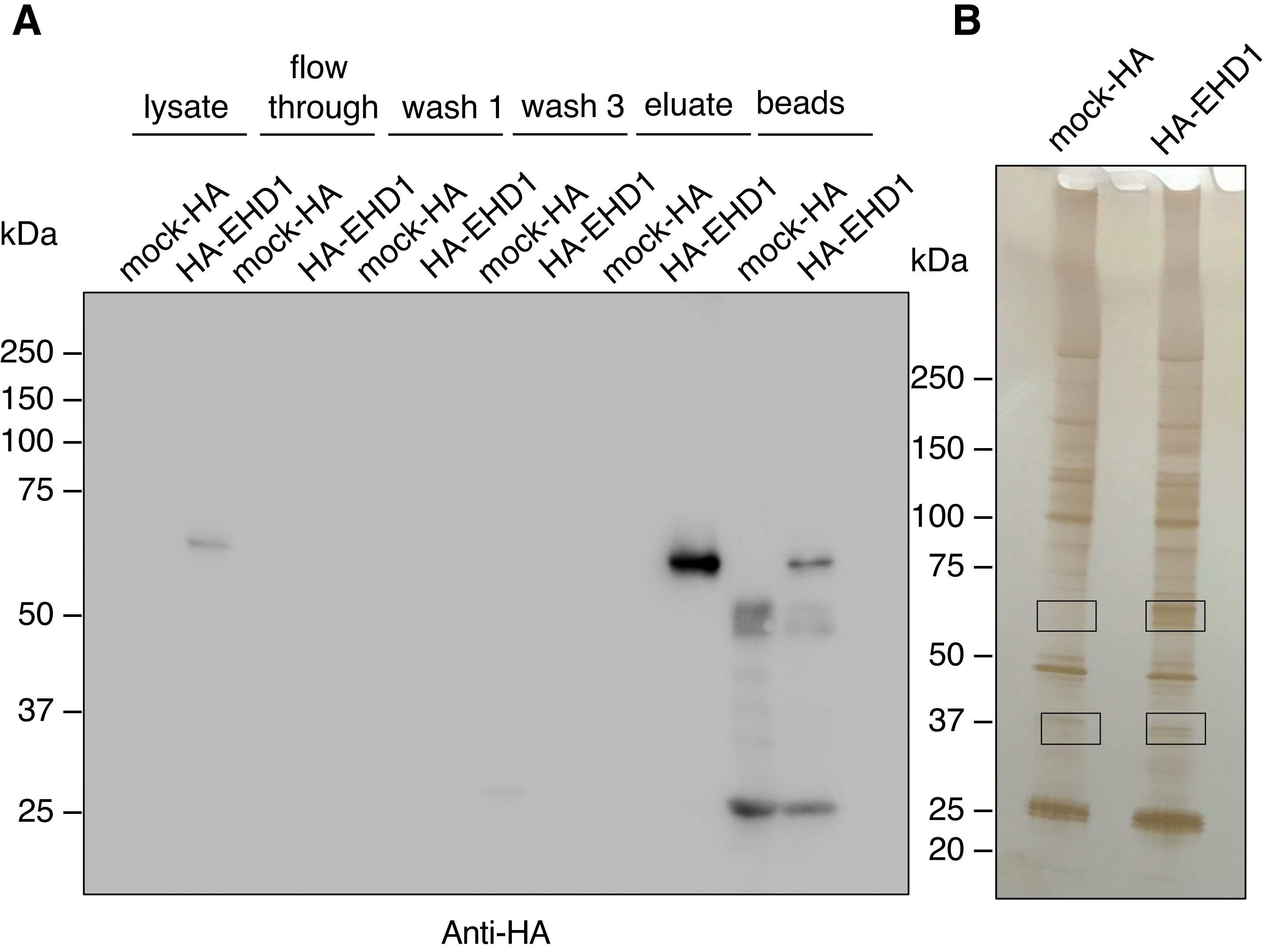
